# Molecular Mechanism of Regulation of RhoA GTPase by Phosphorylation of RhoGDI

**DOI:** 10.1101/2023.05.10.540210

**Authors:** Krishnendu Sinha, Amit Kumawat, Hyunbum Jang, Ruth Nussinov, Suman Chakrabarty

## Abstract

Rho-specific guanine dissociation inhibitors (RhoGDIs) play a crucial role in the regulation of Rho family GTPases. They act as negative regulators that prevent the activation of Rho GTPases by forming complexes with the inactive GDP-bound state of GTPase. Release of Rho GTPase from the RhoGDI-bound complex is necessary for Rho GTPase activation. Biochemical studies provide evidence of a “phosphorylation code”, where phosphorylation of some specific residues of RhoGDI selectively releases its GTPase partner (RhoA, Rac1, Cdc42 etc.). This work attempts to understand the molecular mechanism behind this phosphorylation code. Using several microseconds long atomistic molecular dynamics (MD) simulations of the wild-type and phosphorylated states of the RhoA–RhoGDI complex, we propose a molecular-interaction-based mechanistic model for the dissociation of the complex. Phosphorylation induces major structural changes, particularly in the positively charged polybasic region (PBR) of RhoA and the negatively charged N-terminal region of RhoGDI that contribute most to the binding affinity. MM-PBSA binding free energy calculations show a significant weakening of interaction on phosphorylation at the RhoA-specific site of RhoGDI. In contrast, phosphorylation at a Rac1-specific site leads to the strengthening of the interaction confirming the presence of a phosphorylation code. RhoA-specific phosphorylation leads to a reduction in the number of contacts between the PBR of RhoA and the N-terminal region of RhoGDI, which manifests reduction of the binding affinity. Using hydrogen bond occupancy analysis and energetic perturbation network, we propose a mechanistic model for the allosteric response, i.e., long range signal propagation from the site of phosphorylation to the PBR and buried geranylgeranyl group in the form of rearrangement and rewiring of hydrogen bonds and salt bridges. Our results highlight the crucial role of specific electrostatic interactions in manifestation of the phosphorylation code.

## Introduction

Rho family GTPases, such as RhoA, Rac1, and Cdc42, belong to the Ras superfamily of signaling G-proteins that play a crucial role in several key cellular processes including actin dynamics, cell adhesion, gene transcription (1-5). These proteins share a common sequence and structural features of the G-domain which consists of Switch I and Switch II regions involved in the nucleotide exchange (6-8). Like all other G-proteins, these proteins function as molecular switches that regulate cellular functions by using a simple biochemical strategy of switching between an active guanosine triphosphate (GTP)-bound state and an inactive guanosine diphosphate (GDP)-bound state (8,9). Rho GTPases in their activated state bind to various downstream effectors and regulate processes such as cytoskeletal rearrangement and gene transcription. Several structural and mutational analyses have revealed multiple mechanisms behind the selectivity and specificity between effectors and Rho GTPases (10,11).

Three major proteins control the Rho family GTPases signaling cycle: (i) a GTPase activating protein (GAP), (ii) a guanine nucleotide exchange factor (GEF), and (iii) a Rho-specific guanine nucleotide dissociation inhibitor (RhoGDI) (10). GAP can bind to an activated GTP-bound Rho GTPase and enhance the GTP hydrolysis with the result of terminating the signaling event. GEF activates a monomeric Rho GTPase by stimulating the release of GDP to allow the binding of GTP. RhoGDI binds to a GDP-bound “off” state Rho GTPase and prevents the conversion of the “off” state to the “on’’ state (11-15). Also, it prevents the Rho GTPases from localizing at the membrane which is the place of their action.

RhoGDIs play a critical role in the regulation of Rho family GTPases (16). As described by Garcia-Mata *et al*, RhoGDIs act in the background like an ‘invisible hand’ regulating the level of activated/deactivated Rho GTPases in the cell (17). The population of Rho GTPases at the membrane is increased when RhoGDI is absent. Rho GTPases anchor in the cellular membrane via a prenyl moiety that attaches to the C-terminal cysteine residue and is inserted into the lipid bilayer (18-20). When the RhoGDI binds to this prenylated group, a cytosolic RhoGDI complex with GDP-bound Rho GTPase is formed, which regulates the cytoplasmic pool of each Rho GTPase. Studies using X-ray crystallography have revealed that there are two main regions of the interaction between RhoA and RhoGDI (21-23): (i) the N-terminal region of RhoGDI, which folds into a helix-turn-helix (HTH) motif, interacts with the Switch I and II regions of RhoA, resulting in inhibition of GDP dissociation, and (ii) the C-terminal region of RhoGDI, which folds into a β-sandwich, an immunoglobin-like fold with a hydrophobic pocket, interacts with the prenyl moiety of RhoA.

A complex formation of Rho GTPase with RhoGDI makes the GTPase biologically inert, so activation of Rho GTPase by GEF requires release of Rho GTPase from the complex (24). Experimental studies suggest phosphorylation of RhoGDI as a key post-translational modification for dissociation of Rho GTPase from the complex (25-27). The phosphorylation of RhoGDI is highly specific (supporting information Table S1). For instance, phosphorylation at Ser101 and Ser174 by p21-activated kinase 1 (PAK1) leads to the release of Rac1 (but not RhoA) (24,26). On the other hand, phosphorylation of Ser34 (28) or Ser96 (29) by protein kinase Cα (PKCα) selectively releases RhoA (but not Rac1 or Cdc42). It is speculated that “a unique phosphorylation code” may be at play that controls the release of a specific Rho GTPase from the complex. It is hypothesized that phosphorylation of Ser34 disrupts its interaction with Arg68 of RhoA (for Rac1 or Cdc42, it is Arg66), leading to dissociation (30-32). But this residue is conserved in most of the GTPases, hence the exact mechanism behind the specificity is unclear.

Several NMR studies have shown that two N-terminal regions of RhoGDI (residues 9-20 and 36-58) are highly disordered in absence of Rho GTPases (33-36). The later N-terminal region (residues 36-58) folds into a HTH conformation in presence of Rho GTPases. This HTH conformation is stabilized by the formation of several hydrogen bonds (H-bonds) between conserved residues of Switch I (Thr37 and Val38) of RhoA (Thr35 and Val36 for Rac1 and Cdc42) and RhoGDI (Asp45 and Ser47) (37). The N-terminal region (residues 9-20) can exist either in helix or random coil conformations (38). Recent crystal structures show the presence of helix (residues10-15) in RhoA (PDB ID: 4F38)(39) and Cdc42 (PDB ID: 1DOA)(22), whereas for Rac1 it exists as a random coil (PDB ID: 1HH4) (23).

In this present work, we have employed several microseconds long atomistic molecular dynamics (MD) simulations to understand the conformational dynamics and energetics of the RhoA–RhoGDI complex both in the wild-type and two different phosphorylated states of RhoGDI. We first attempt to establish the presence of phosphorylation code by showing that RhoA-specifc phosphorylation at Ser34 leads to reduction in binding energy between RhoA and RhoGDI, whereas Rac1-specific phosphorylation at Ser101/Ser174 does not. Subsequently, we present a thorough structural and energetic analysis to establish a mechanistic picture of how the effect of RhoA-specific phosphorylation propagates to a distal site allosterically to cause the reduction in binding affinity. Essentially, the goal of this work is to establish the molecular thermodynamic origin of the specificity of the phosphorylation code in the RhoA–RhoGDI interaction.

## Computational Methods

### Structure Preparation

For modeling and simulation, we used the crystal structure of RhoA–RhoGDI complex (PDB ID: 4F38) (Figure 1). Using the PyMol program (40), we replaced the GNP molecule present in the crystal structure with GDP. The pdbfixer utility of OpenMM package was used to model the missing residues and atoms (41). We utilized the PyTMs plugin of PyMol to prepare the phosphorylated system. For our study, we constructed three different systems: (i) WT (wild type): RhoA-GDP in complex with wild-type RhoGDI, (ii) SP34: RhoA-GDP in complex with phosphorylated Ser34 (pS34) RhoGDI, and (iii) SP101/174: RhoA-GDP in complex with phosphorylated Ser101/Ser174 (pS101/pS174) RhoGDI.

**Figure 1:**
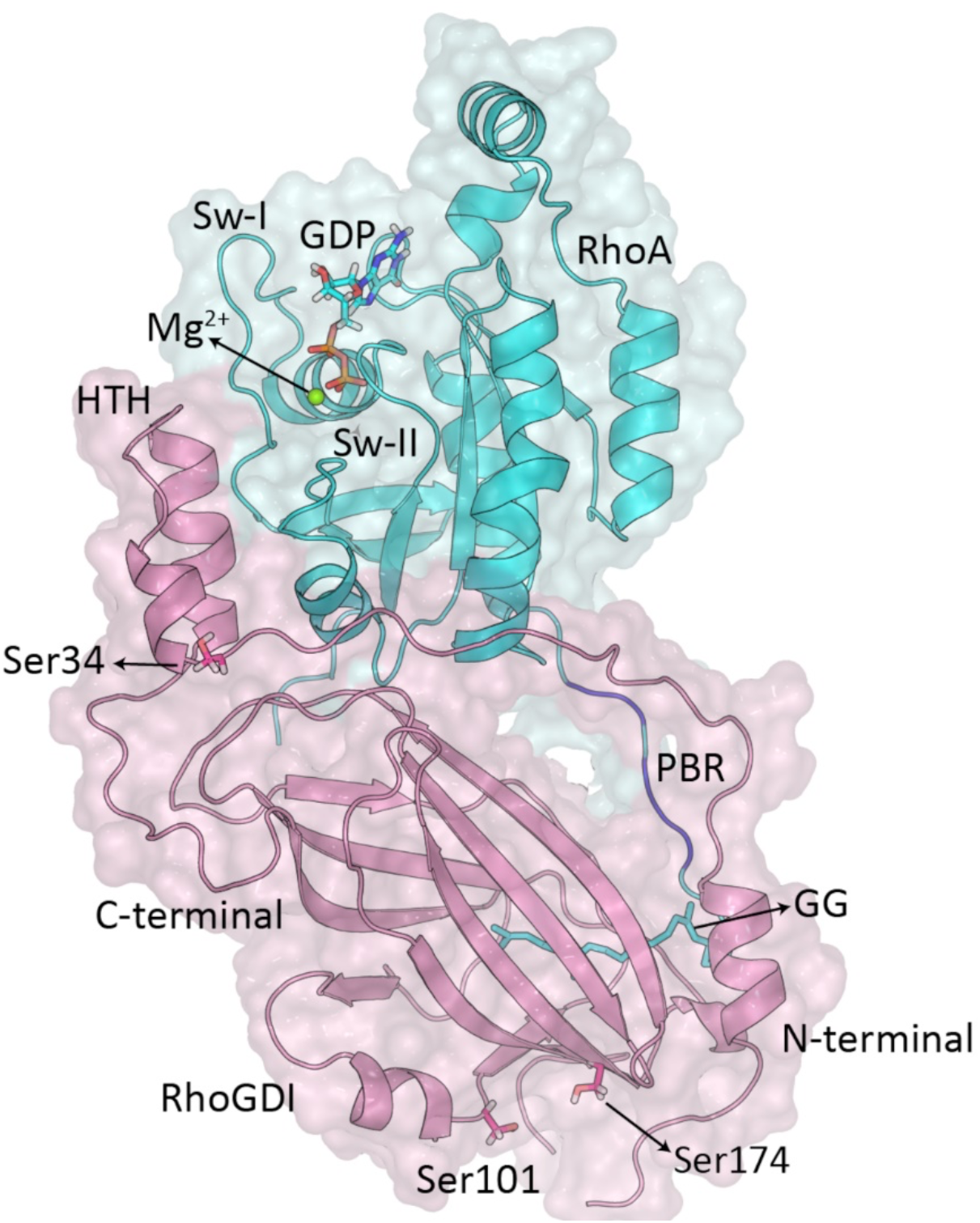
Crystal structure of RhoA–RhoGDI complex. RhoA (residues 1-190) is shown in cyan color, and RhoGDI (residues 1-204) is shown in pink color. The PBR region of RhoA is shown in blue. The sites of phosphorylation (Ser34, Ser101 and Ser174), GDP, and the geranylgeranyl (GG) moiety are shown in stick representation. Mg^2+^ is shown as a green sphere.

### Molecular Dynamics (MD) Simulations

We have performed 4 independent 1 microsecond long MD simulations of these three systems. All the simulations were performed using GROMACS 2019.6 software (42). Charmm36 force field was used along with the TIP3P water model (43,44). The energy minimization of the solvated proteins was accomplished using the steepest descent algorithm. A modified Berendsen thermostat (45) at 300 K and Parrinello-Rahman barostat (46) at 1 bar were utilized in NPT ensemble for the production runs. Long-range electrostatic interactions were calculated with the particle mesh Ewald (PME) summation method (47) with a grid spacing of 0.16 nm and fourth-order cubic interpolation. The cut-off distance used for short-range electrostatic and van der Waals (vdW) interactions was 1 nm. All covalent bonds were constrained using the LINCS algorithm (48). The integration time step was set to 2 fs.

### Electrostatic Interaction Energy Calculations

Phosphorylation means the addition of an extra negative charge to the system. We are interested in learning how the introduction of a single negative charge affects structural modification at a distant point. We have established in our earlier work that electrotatic interaction energy can be a sensitive reporter of subtle structural changes including long range allosteric effects (49,50). Using a similar approach, here we have computed the perturbation/change in the average interaction energy for each residue (Δ⟨*E*_*i*_⟩) upon phosphorylation, i.e. we have compared between the wild-type 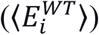 and phosphorylated 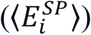 states of the complex. The change in average interaction energy of *i*-th residue, Δ⟨*E*_*i*_⟩, is given by

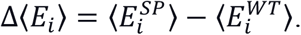

We have analyzed the pairwise changes in interaction energy and H-bond occupancy to determine how the signal propagates from the site of perturbation (phosphorylation) to the protein-protein interaction interface. The change in average pair-wise interaction energy of *i*-th and *j*-th residue pair is given by

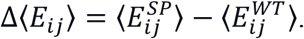

### Binding Free Energy Calculations

The binding free energy between RhoA and RhoGDI was calculated using the molecular mechanics Poisson-Boltzmann surface area (MM-PBSA) method. The GROMACS tool, g_mmpbsa software (51) was used to calculate the binding free energy. The binding free energy is estimated as the difference between the free energy of the RhoA–RhoGDI complex and the free energies of the unbound components (RhoA and RhoGDI). The total binding free energy, Δ*G*_*bind*_, is calculated as a sum of the gas-phase contribution from the molecular mechanics energy, Δ*E*_*MM*_, the solvation energy associated with the transition of the ligand from bulk water to the binding site, Δ*G*_*sol*_, and the change in conformational entropy related to the binding of the ligand, −*T*Δ*S*.

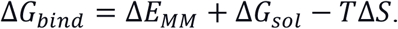

In our work, we did not consider the entropic component of the binding free energy. Δ*E*_*MM*_ is calculated based on molecular mechanics model of the underlying force field. It consists of different components, the electrostatic interaction, Δ*E*_*elec*_, and the vdW interaction, Δ*E*_*vdW*_. The polar contribution to the solvation free energy, Δ*G*_*PB*_, is estimated using the Poisson-Boltzmann implicit solvent model, and the non-polar contribution, Δ*G*_*SA*_, is determined using the solvent accessible surface area. The set of parameters used for MM-PBSA calculation is given in supporting information (Table S2).

### Principal Component Analysis (PCA)

PCA is a powerful dimensionality reduction technique that enables us to project the higher dimensional MD simulation data to a lower dimensional representation for easier visualization and interpretation of the conformational landscape sampled by the proteins. The basic idea of PCA is to diagonalize the covariance matrix of the higher dimensional dataset and use the eigenvectors corresponding to leading eigenvalues to construct the lower dimensional representation. In this work, we have considered projections along the top two eigenvectors, i.e., PC1 and PC2.

### Coordination Number (CN) as a collective variable

The collective variable coordination number (CN) represents the number of contacts formed between two groups of atoms (A and B) defined as:

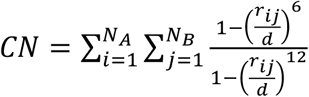

where *i* and *j* are the indices of a set of selected atoms, and *N*_*A*_ and *N*_*B*_ represent the number of atoms present in group A and group B. *r*_*ij*_ is the distance between the *i*-th and *j*-th atoms. The cutoff distance, *d*, was chosen as 3.0 Å in order to include the distance criteria of the H-bond. Thus, the coordination number provides an estimate of the number of H-bonds that are formed between selected groups. The coordination number was calculated using the PLUMED 2.5 software (52).

## Results and Discussion

### MM-PBSA Analysis of Binding Free Energy Between RhoA and RhoGDI Proteins

Motivated by the idea of the presence of a phosphorylation code as discussed earlier, we have compared the binding free energy between RhoA and RhoGDI across three different systems: (i) WT (wild type): RhoA-GDP in complex with wild-type RhoGDI, (ii) SP34: RhoA-GDP in complex with phosphorylated Ser34 (pS34) RhoGDI, and (iii) SP101/174: RhoA-GDP in complex with phosphorylated Ser101/Ser174 (pS101/pS174) RhoGDI. On the RhoA-specific phosphorylation (SP34 system), the binding of RhoA to RhoGDI is expected to become weaker, but not for the Rac1-specific phosphorylation (SP101/174 system).

A reduction in binding affinity (less negative values) signifies a weaker interaction between RhoA and RhoGDI. Figure 2a shows the probability distribution of binding free energy for the different systems. For Ser34 phosphorylation, the binding free energy increases (less negative), whereas for Ser101 and Ser174 phosphorylation, the binding free energy slightly decreases (more negative) as compared to the WT system. Residue-wise decomposition of binding free energy for the WT system shows two major binding regions (Figure 2b): (i) Switch II of RhoA and HTH motif of RhoGDI, and (ii) PBR of RhoA and N-terminal region (residues 9-25) of RhoGDI. The residues with a large contribution to overall binding free energy are shown in blue color (Figure 2c).

**Figure 2.**
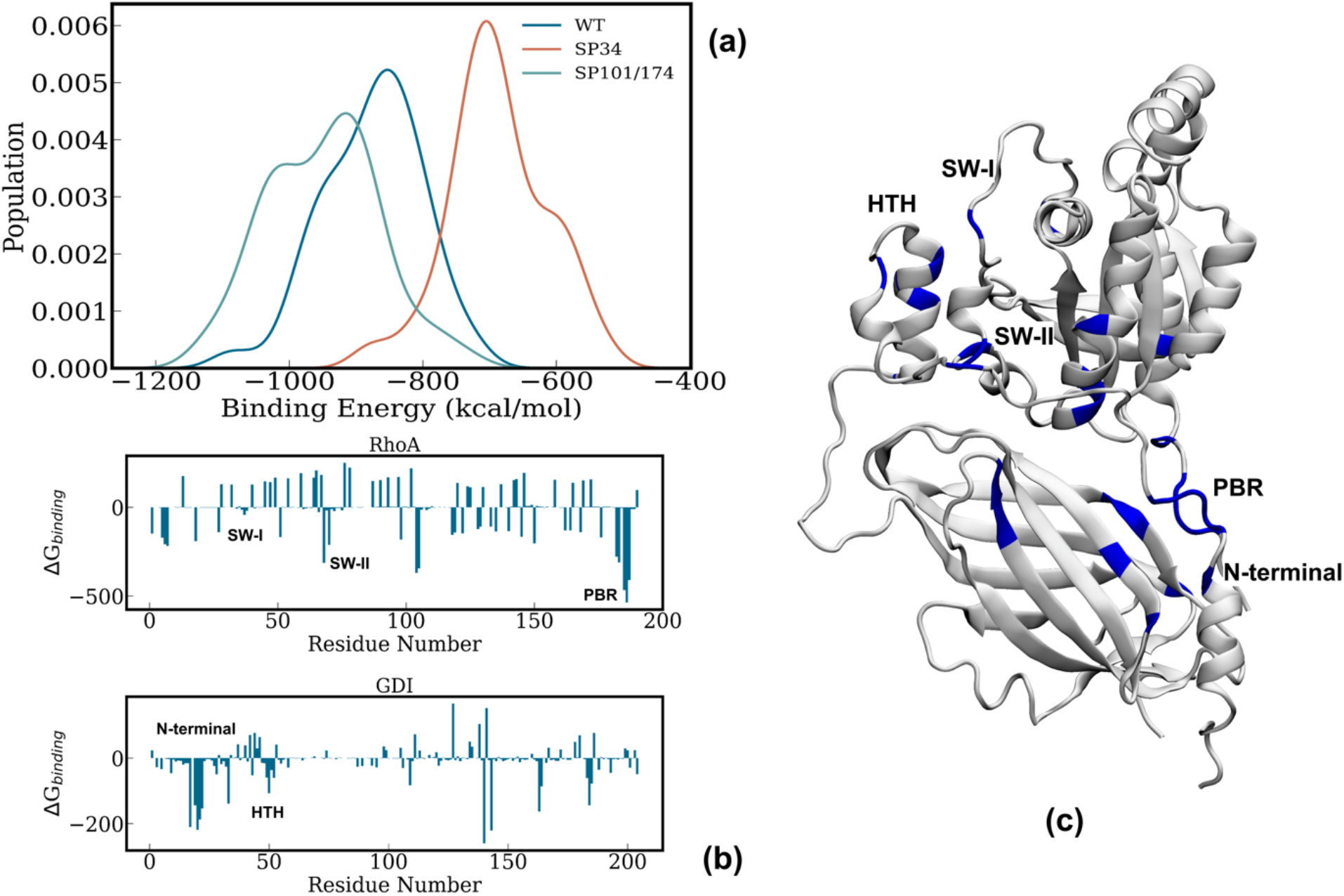
Analysis of binding free energy of the RhoA–RhoGDI complex using MM-PBSA. (a) Comparison of the binding free energy distribution between WT, SP34, and SP101/174 systems. (b) Residue-wise decomposition of binding free energy in the WT system for RhoA and RhoGDI proteins. (c) Structure of WT system, where the residues with favorable contributions to binding free energy (Δ⟨*E*_*i*_⟩ < 0) are highlighted in blue.

Figure 3a shows the residue-wise change in binding free energy for the phosphorylated systems. It can be seen that the major difference between Ser34 and Ser101/Ser174 phosphorylation arises from the interaction between RhoA PBR and RhoGDI N-terminal region. For the SP34 system, the interaction of PBR with the N-terminal region is weakened, while it is strengthened for SP101/174 system. Visual inspection of the structural superposition in Figure 3b shows that the distance between PBR and N-terminal region becomes higher for the SP34 system, while it is decreased for the SP101/174 system compared to the WT. As a result, the binding free energy is increased for the SP34 system, while it is decreased for the SP101/174 system. This is in accordance with previous experimental studies (28,29) and also provides evidence for the existence of a phosphorylation code.

**Figure 3.**
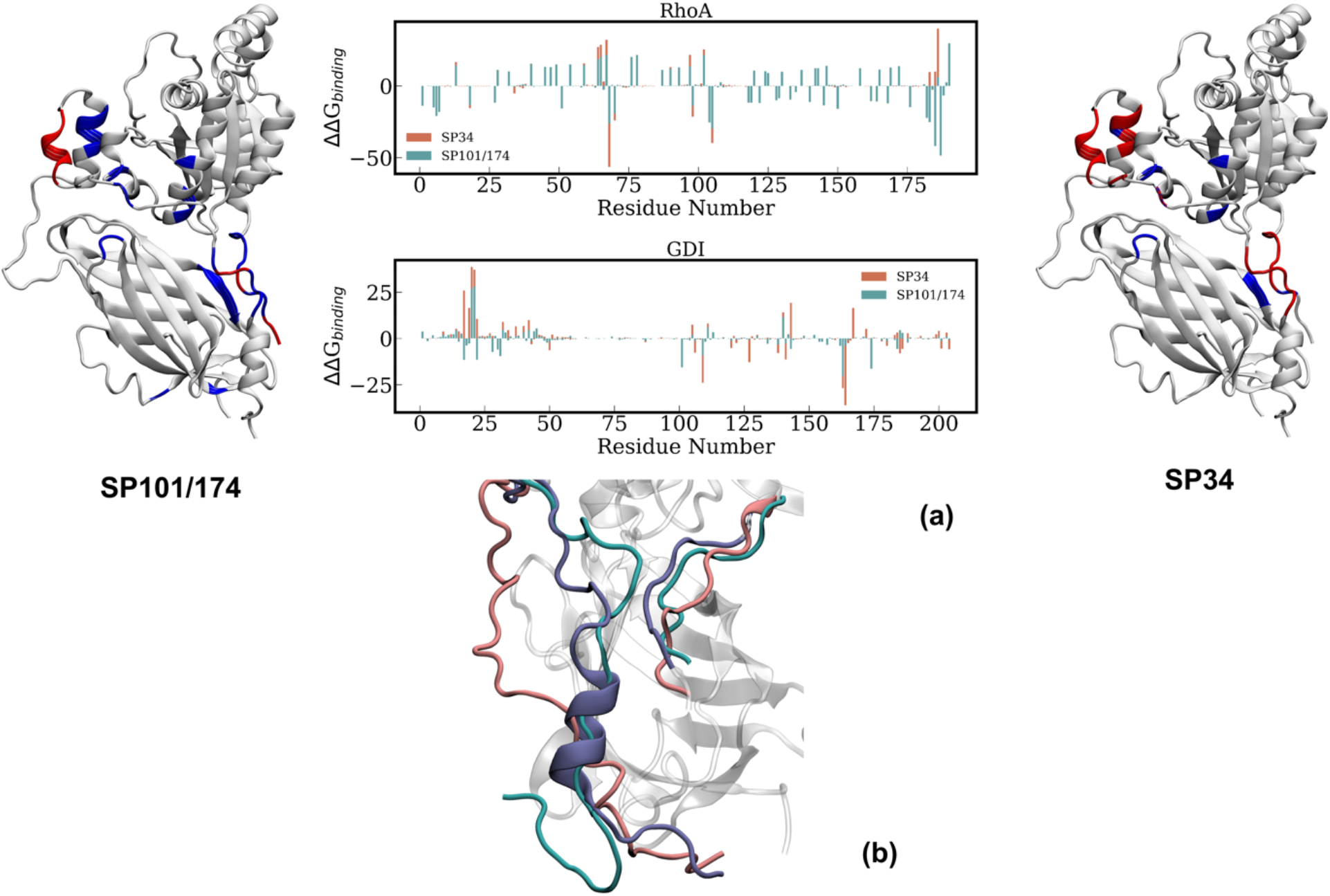
a) Residue-wise contribution to the change in binding free energy. Residues with unfavorable (>0) contribution is shown in red and favorable (<0) contribution is shown in blue color. b) Position of PBR and N-terminal region for WT, SP34 and SP101/174 systems. WT is shown in blue, SP34 is shown in pink, and SP101/174 is shown in teal color.

Now that we have clearly established that phosphorylation at Ser34 leads to reduction in the RhoA–RhoGDI interaction, the subsequent sections will focus specifically on the SP34 system to unravel the molecular mechanism behind how the effect of phosphorylation manifests into reduction of the binding affinity.

### Structural Changes Due to Ser34 Phosphorylation

It is expected that the reduction in binding affinity between RhoA and RhoGDI after phosphorylation should be associated with some structural changes that affect the protein-protein interaction (PPI) interface. Root-mean-square deviation (RMSD) was computed for both RhoA and RhoGDI with references to the crystal structure to monitor such structural deviations, if any. The RMSD remains more or less stable for RhoA, whereas for RhoGDI it changes considerably around 600 ns (Figure 4a). By visual inspection, we have observed three significant structural changes after phosphorylation. These are (i) melting of the N-terminal helix, (ii) change in the conformation of the N-terminal loop, and (iii) outward movement of the geranylgeranyl group of RhoA from the hydrophobic cavity in the C-terminal β-sandwich of RhoGDI.

**Figure 4.**
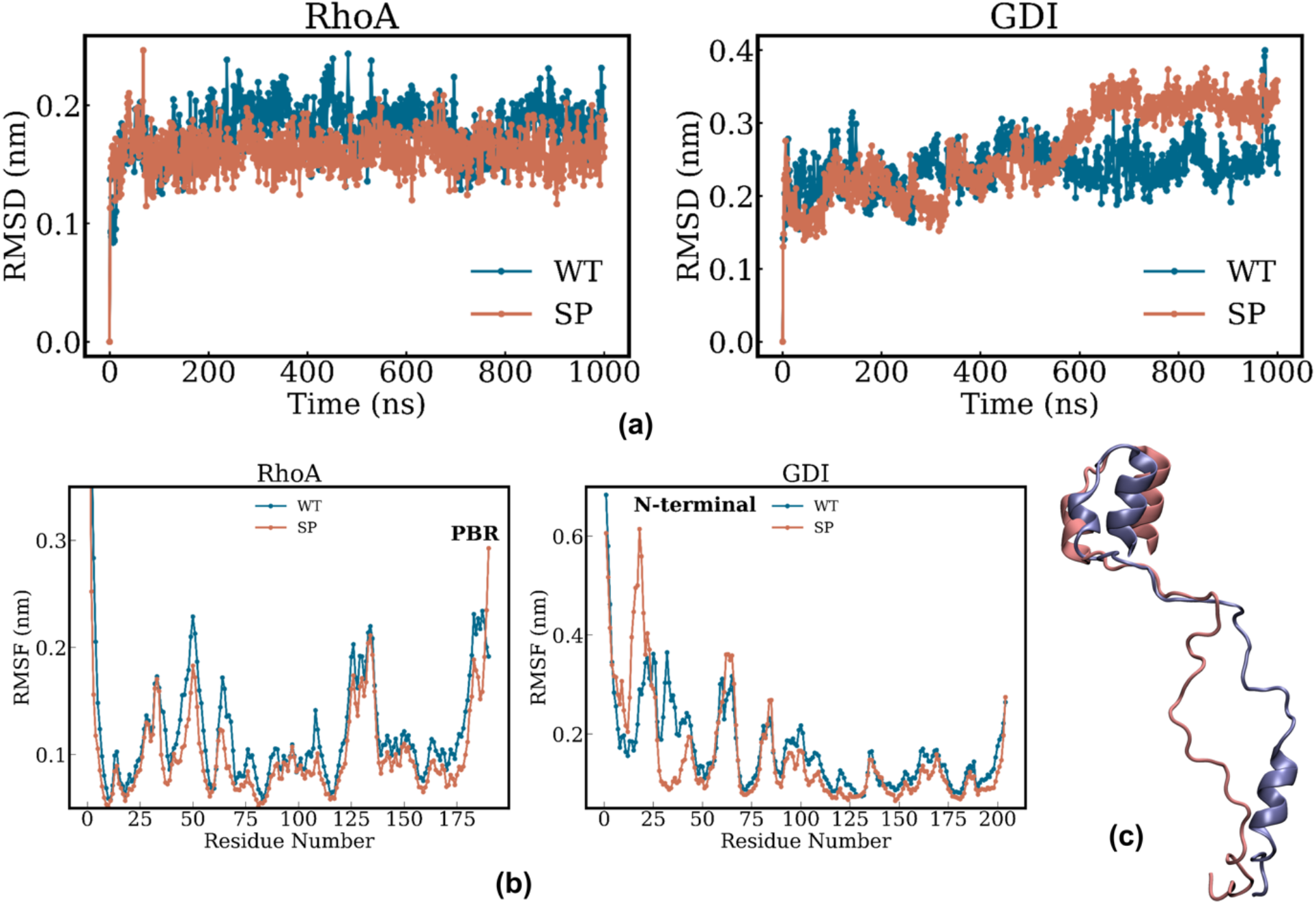
(a) Time series of RMSD for WT and SP34 systems. (b) Residue-wise RMSF for RhoA and RhoGDI separately. (c) Superimposed structures of the N-terminal region for WT and SP34 systems.

We have also compared the residue-wise root-mean-square fluctuation (RMSF) between the wild-type (WT) and phosphorylated (SP) systems to investigate the effect of phosphorylation on the local dynamics/flexibility (Figure 4b). The major changes in residue-wise RMSF are observed in the PBR of RhoA (residues 182-187) and the N-terminal region of RhoGDI (residues 1-40) (Figure 4c). Also, there are some small differences in the Switch I (residues 28-42) and Switch II (residues 61-81) loops of RhoA. Interestingly, these switch loops are known to be crucial for the function of Rho GTPases and they can exhibit nucleotide dependent conformational heterogeneity as reported earlier (7). The largest change in RMSF was observed in the N-terminal region of RhoGDI (Figure 4c).

In order to fully characterize the phosphorylation induced modulation in the conformational ensemble and dynamics, we have performed principal component analysis (PCA) on the combined WT and SP34 trajectories (Figure 5). It is evident that the major difference in protein conformation is captured by the 1st principal component (PC1) followed by the 2nd (PC2) (Figure 5a). These two dynamical modes are visualized by the porcupine plots of the corresponding eigenvectors. For SP34, PC1 eigenvectors show an outward movement of the N-terminal region of RhoGDI, which pulls the PBR of RhoA (Figure 5b). In PC2, an upward movement of the C-terminal β-sandwich of RhoGDI is observed (Figure 5c). PC2 eigenvectors also show the movement of the N-terminus of RhoGDI towards the C-terminal β-sandwich. The outward movement of the N-terminal region pulls the PBR, as the PBR moves away from the C-terminal β-sandwich, which weakens the geranylgeranyl–cavity interaction, ultimately leading to the dissociation of the complex.

**Figure 5.**
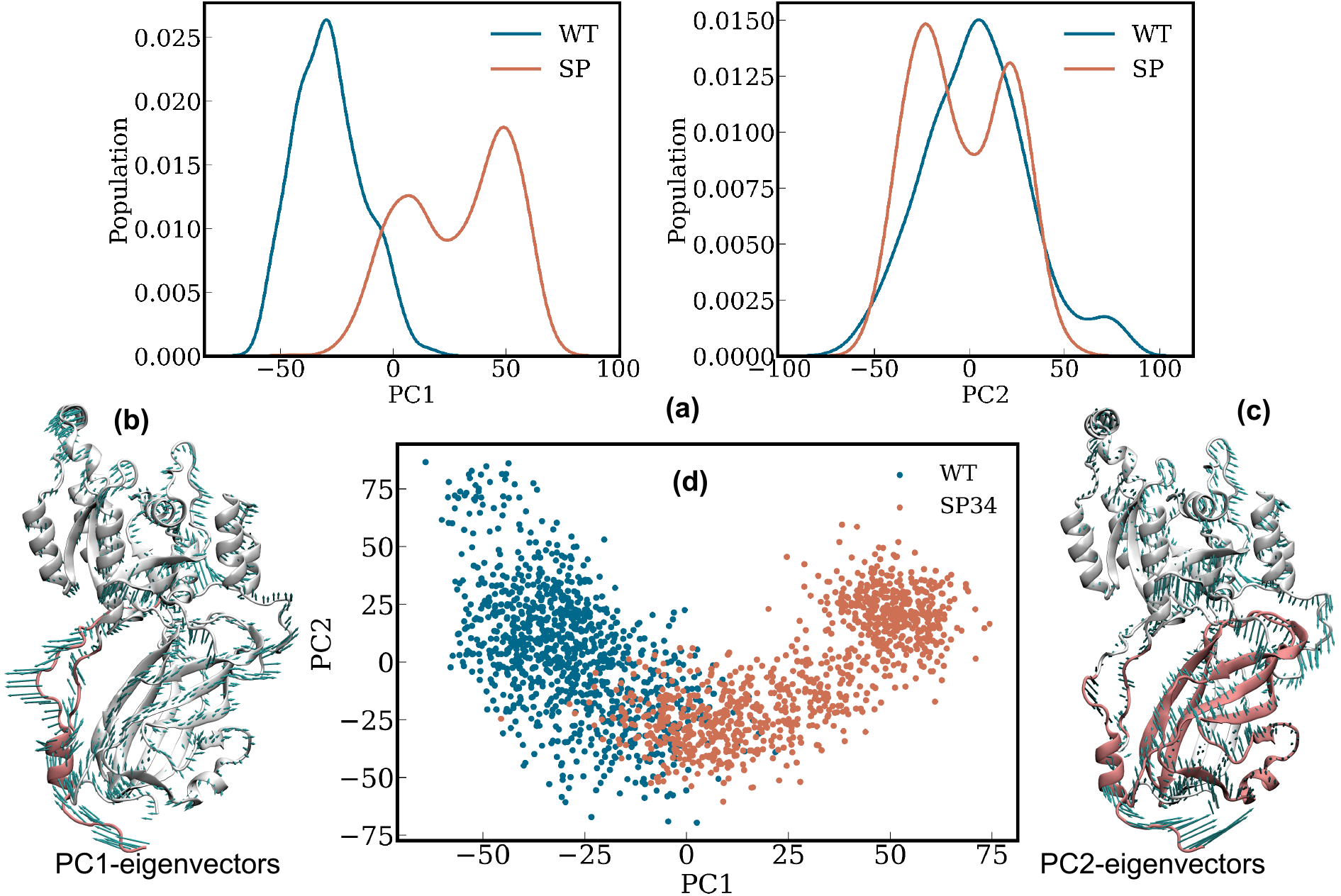
Principal component analysis (PCA): (a) Distributions of PC1 and PC2. Projections of eigenvectors on protein structure for (b) PC1 and (c) PC2. (d) The projection of the first two principal components, PC1 and PC2, for WT and SP34 systems.

As mentioned earlier, two major structural rearrangements observed upon phosphorylation are: (i) the N-terminal region of RhoA moving away from the PBR region of GDI, and (ii) loosening (pulling out) of the geranylgeranyl hydrophobic tail from the GDI cavity. We have further characterized these changes in terms of more localized structural parameters. First, we monitor the number of contacts or coordination number (CN) between (a) the N-terminal and PBR, and (b) the N-terminal and C-terminal of GDI (Figure 6a). The time evolution of these contacts clearly show that for the phosphorylated SP34 system, the CN between the N-terminal region of RhoGDI and PBR of RhoA decreases significantly, while the contacts between the N-terminal and C-terminal regions of RhoGDI increase (Figure 6a). This indicates that the N-terminal region moves away from the PBR but moves towards its C-terminal region as shown strucrally in Figure 6b. A decrease in the CN between the N-terminal region and PBR leads to decreased interaction between RhoA and RhoGDI, leading to a specific release of RhoA from RhoGDI.

**Figure 6.**
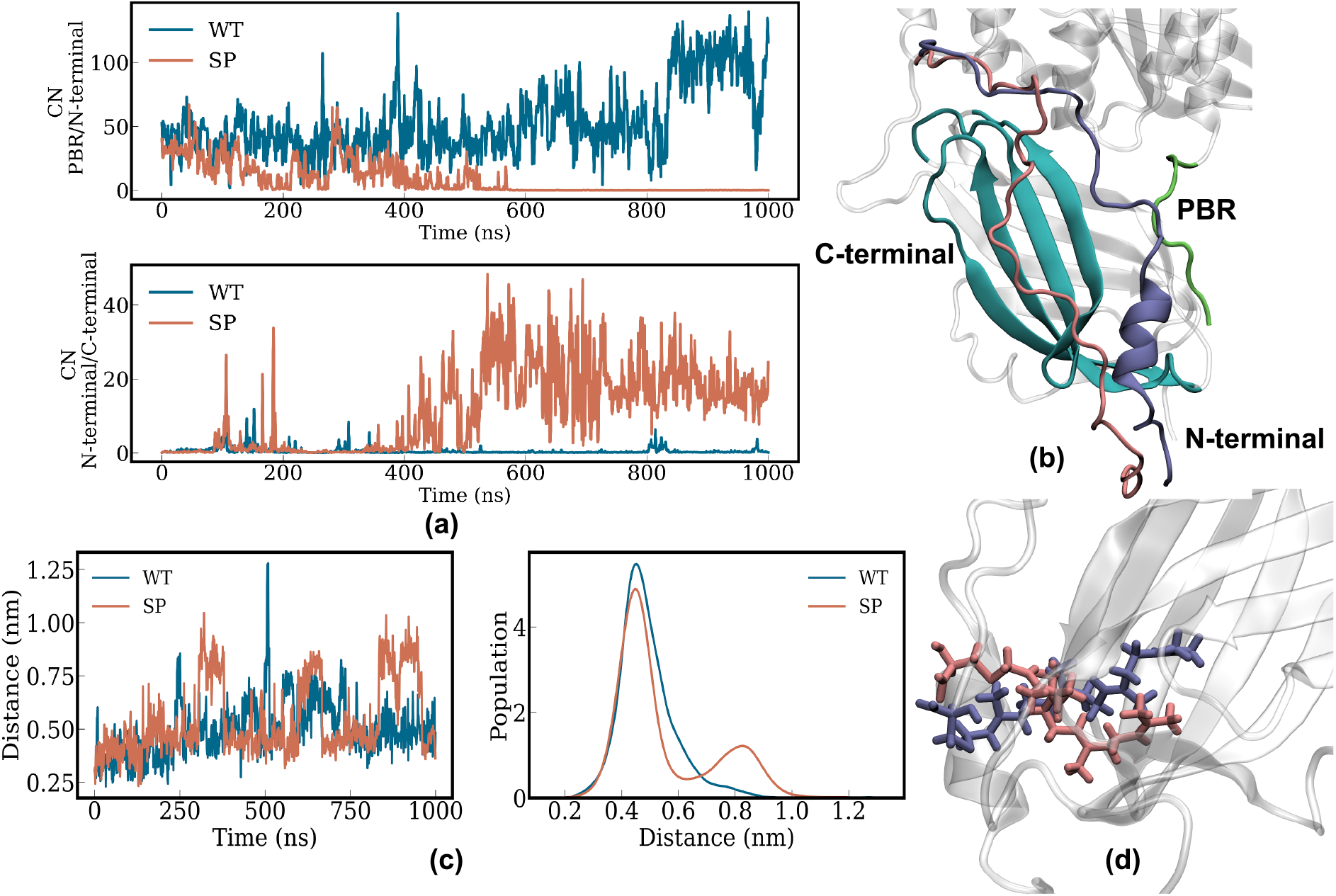
Coordination number (CN) analysis. (a) Time series of the CNs between the N-terminal region of RhoGDI and RhoA PBR (upper panel), and between the N-terminal and C-terminal regions of RhoGDI (lower panel). (b) Snapshot representing the movement of N-terminal region of RhoGDI away from RhoA PBR. (c) Time series of the distance (left panel) and distribution of the distance (right panel) between the geranylgeranyl group and hydrophobic cavity. (d) Snapshot highlighting the outward movement of the geranylgeranyl group.

In order to monitor the “pulling out” motion of the geranylgeranyl tail, we calculated the distance between the geranylgeranyl group of RhoA and the center of mass (COM) of the hydrophobic cavity of RhoGDI (Figure 6c). This gives us an idea of how much the prenyl moiety is moving away from the hydrophobic cavity. It is evident from the increased distance that after phosphorylation that the geranylgeranyl group of RhoA tends to come out of the hydrophobic cavity of RhoGDI (Figure 6d). We observe that the orientation of the N-terminal region also changed significantly due to phosphorylation.

### Propagation of Signal and Mechanism of Dissociation

The addition of extra negative charges due to phosphorylation creates a local perturbation to the electrostatic interaction network. Such redistribution of electrostatic interactions in terms of rewiring of hydrogen bonds and salt bridges in a domino-like fashion can be a dominant mechanism of allostery as we have demonstrated earlier (49,50,53). In addition, electrostatic interaction can be a very sensitive reporter of rather subtle changes in the conformational ensemble (49). For this reason, we track the residue-wise changes in average interaction energy to understand how the energetic perturbation created at the site of phosphorylation propagates to the PPI interface and weakens it. The change in residue-wise average interaction energy is defined as 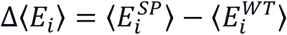, where 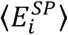 and 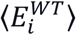 are the average non-bonded interaction energy of the *i*^th^ residue with its environment (protein, water and ions).

From Figure 7a, it can be seen that the largest change is at the site of phosphorylation, i.e., Ser34, as expected. But in addition, significant change in the interaction energy is visible near the PBR and N-terminal regions involved in stabilizing the PPI. In Figure 7b, we visualize the distribution of these residues with significant perturbation in interaction energy. The Cα atoms of each residue is shown as a sphere, where the size of the sphere represents the magnitude of Δ⟨*E*_*i*_⟩. Blue and red spheres indicate negative and positive signs of Δ⟨*E*_*i*_⟩, respectively. It is evident that the largest changes are observed near the phosphorylation site and the polybasic region of RhoA with a few residues being visible in the intermediate regions. Figure 7b clearly highlights the dramatic long range (allosteric) perturbation created by the phosphorylation that manifests in the substantial structural and energetic rearrangement in the PBR and N-terminal region as mentioned earlier.

**Figure 7.**
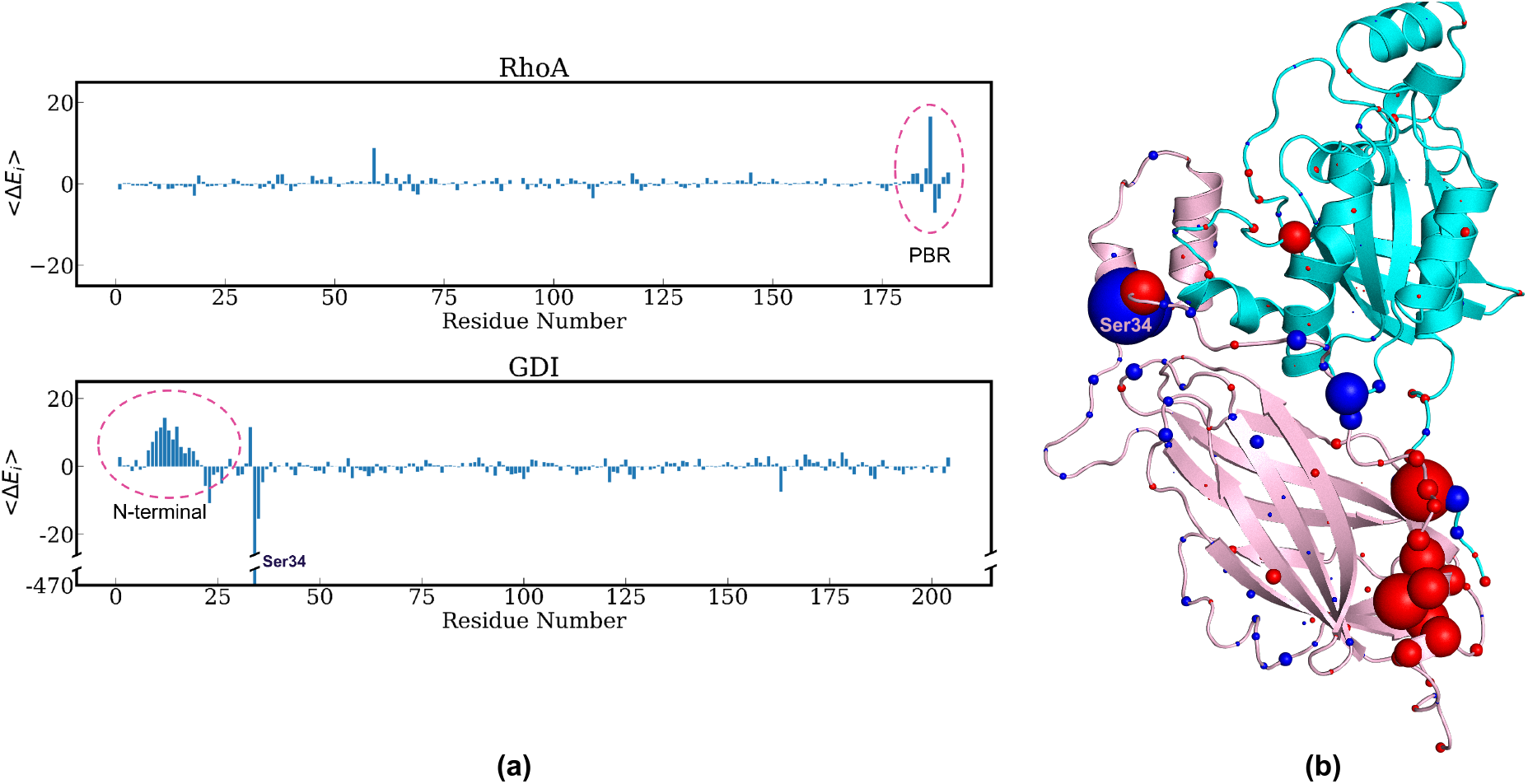
(a) Residue-wise change in the interaction energy (Δ*E*_*i*_) for RhoA and GDI. (b) CA atom of each residue shown on protein structure, the size of the sphere indicate the absolute value of Δ*E*_*i*_ and color of the sphere indicates the sign of Δ*E*_*i*_. Blue color indicates it is negative and red color indicates it is positive. Largest change is observed at the site of phosphorylation.

The next obvious question to address would be how does the signal or energetic perturbation created by the phosphorylation at Ser34 propagate over such a long distance (>3.5nm) from the site of phosphorylation to PBR region of RhoA? In order to track the pathway of the propagation of this energetic perturbation, we have interrogated the rewiring of the underlying interaction energy (Δ⟨*E*_*ij*_⟩) network. Essentially, we compute the residue pairwise average interaction energy (⟨*E*_*ij*_⟩) and how much that changes upon phosphorylation: Δ⟨*E*_*ij*_⟩. Then we can easily build a network of residue-pairs with significant changes in average interaction energy and track how the perturbation propagates as shown in Figure 8. Blue and red lines here indicate pair-wise interactions that became stronger (Δ⟨*E*_*ij*_⟩ < 0) and weaker (Δ⟨*E*_*ij*_⟩ > 0), respectively, upon phosphorylation. The Δ⟨*E*_*ij*_⟩ value for each residue pair is reported in supporting information (Table S3). We have also observed that most of these pairs show a significant change in the hydrogen bond occupancy Δ⟨*Hb*_*ij*_⟩ (values are reported in Table S4 of supporting information). Several such pairs form a salt bridge either in the WT or SP34 system. Such a network enables us to easily visualize the sequence of molecular events in terms of local rearrangement and rewiring of interactions (particularly, hydrogen bonds and salt bridges) that connect the site of phosphorylation (Ser34) to the distal PBR region.

**Figure 8:**
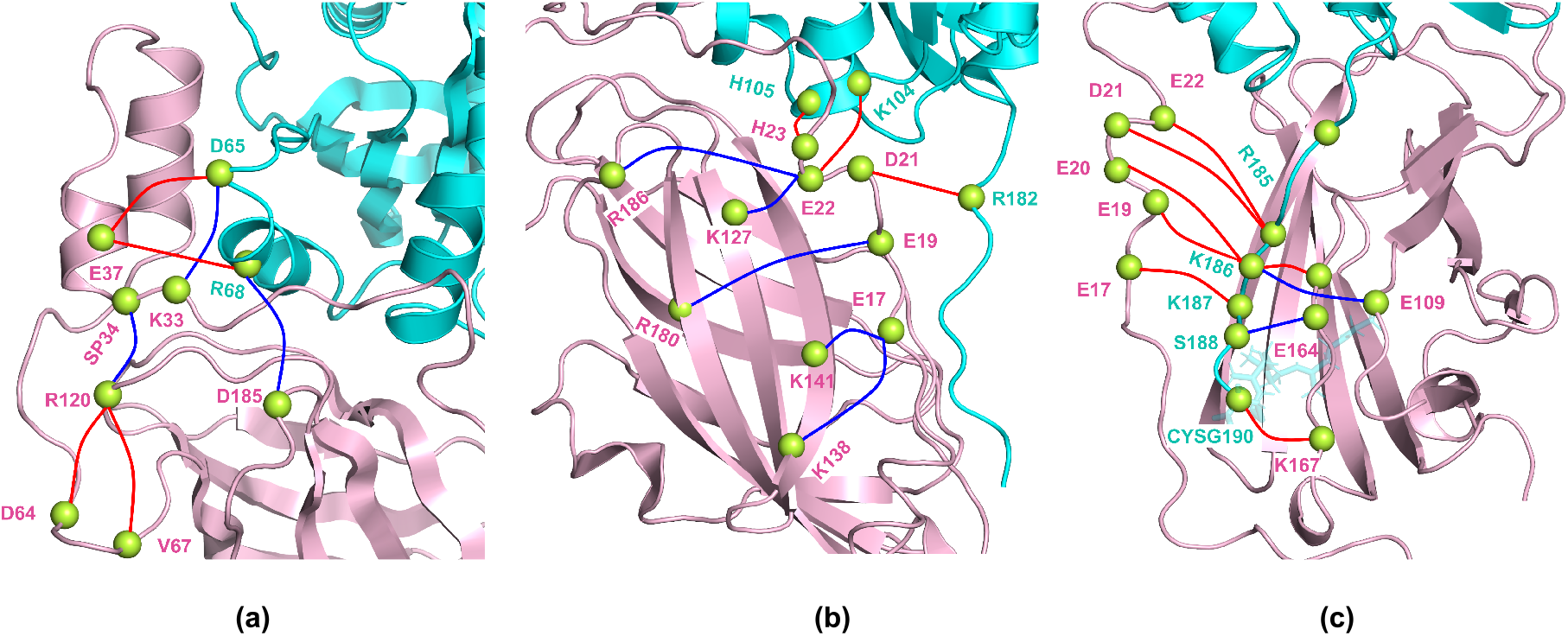
Mechanistic picture of signal propagation from site of phosphorylation (SP34) to PBR region in terms of change in residue pairwise average interaction energy (Δ⟨*E*_*ij*_⟩) for (a) the HTH region, (b) the C-terminal β-sandwich, and (c) the N-terminal region of RhoGDI. Blue and red lines indicate the residue-pairs where the interaction becomes stronger (Δ⟨*E*_*ij*_⟩ < 0) and weaker (Δ⟨*E*_*ij*_⟩ > 0), respectively, upon phosphorylation.

As seen in Figure 8a, after phosphorylation at Ser34, since a negative charge is introduced at pS34, it forms a salt-bridge with Arg120 of RhoGDI. Arg68 of RhoA, which had a favorable interaction with Glu37 earlier, now moves to form an intermolecular H-bond with Asp185 of RhoGDI. As a result, the N-terminal region of RhoGDI becomes free and more flexible. This increased flexibility breaks the intermolecular interactions of Glu22^RhoGDI^ with Lys104^RhoA^ and Arg182^RhoA^ (Figure 8b). The N-terminal region moves towards the C-terminal region of RhoGDI. This conformation is stabilized by a few newly formed intramolecular H-bonds (e.g., Glu22/Arg186, Glu22/Lys127, Glu19/Arg180, Glu17/Lys141, and Glu17/Lys138). It is also interesting to note that the PBR region breaks several hydrogen bonds including a few with the N-terminal region (Figure 8c). This is associated with the PBR region severing its ties with N-terminal and forming a few new contacts with the C-terminal region. A mechanistically important observation is that the salt bridge between CYSG190^RhoA^ and Lys167^RhoGDI^ breaks upon phosphorylation, which is essential for the geranylgeranyl group coming out of the hydrophobic cavity.

An interesting point to note here is that after phosphorylation, the interaction between Ser188^RhoA^ forms stronger interaction with Glu163^RhoGDI^ and Glu164^RhoGDI^. Previously it has been observed that phosphorylation of Ser188^RhoA^ leads to the stabilization of the RhoA-RhoGDI complex (54). This observation is consistent with our results, since phosphorylation of Ser188^RhoA^ (introduction of negative charge) would lead to repulsion with negatively charged Glu163^RhoGDI^ and Glu164^RhoGDI^. According to our results, the formation of Ser188^RhoA^ and Glu164^RhoGDI^ interaction seems to be an important intermediate state towards eventual dissociation of the RhoA–RhoGDI complex.

## Conclusions

In this study, we have established the existence of a “phosphorylation code” in the selective dissociation of RhoA-RhoGDI complex. Our several microseconds long MD simulation trajectories provide a clear microscopic and mechanistic picture of the initial phases of phosphorylation. The picture illustrates structural changes including the weakening of protein-protein interaction only upon phosphorylation at the RhoA-specific site (Ser34). MM-PBSA binding free energy calculations clearly establish that there is a significant increase in binding free energy for the complex formation in the case of RhoA-specific phosphorylation at Ser34, but no major change for phosphorylation at Ser101/Ser174, presumed to be the functional site of phosphorylation to release Rac1 instead of RhoA. We must note here that the binding between RhoA and RhoGDI mainly originates from the strong interaction of the negatively charged N-terminal region of RhoGDI and the positively charged PBR of RhoA. When Ser34 is phosphorylated, the interaction between the HTH region of RhoGDI and the switch region of RhoA is decreased, allowing the N-terminal loop more flexible and moving away from the PBR. The major structural changes are movement of N-terminal region of RhoGDI from PBR of RhoA to the C-terminal region of RhoGDI, and also a “pulling out” motion of the geranylgeranyl group from the hydrophobic cavity of RhoGDI. We have used perturbation in residue-wise (and pair-wise) interaction energy as a reporter of the structural rearrangements that are responsible for propagation of the energetic perturbation created by the phosphorylation. We propose a mechanistic model for the distant signal propagation from the site of phosphorylation to the PBR and buried geranylgeranyl group in the form of rearrangements and rewiring of hydrogen bonds and salt bridges (charge-charge interactions), which demonstrates the crucial role of local and specific electrostatic interactions in the manifestation of observed allosteric response and “phosphorylation code”.

## Acknowledgements

This work has been partially funded by Science and Engineering Research Board (SERB), Govt. of India (project number: ECR/2018/002903). This project has been funded in whole or in part with federal funds from the National Cancer Institute, National Institutes of Health, under contract HHSN261201500003I. The content of this publication does not necessarily reflect the views or policies of the Department of Health and Human Services, nor does mention of trade names, commercial products, or organizations imply endorsement by the U.S. Government. This research was supported in part by the Intramural Research Program of the NIH, National Cancer Institute, Center for Cancer Research. All simulations have been performed using Technical Research Centre (TRC) computing facilities of S. N. Bose National Centre for Basic Sciences, established under the TRC project of Department of Science and Technology (DST), Govt. of India.

## References

1. Bishop, A. L., and A. Hall. 2000. Rho GTPases and their effector proteins. Biochemical Journal. 348(2):241–255, doi: 10.1042/bj3480241, https://doi.org/10.1042/bj3480241.

2. K. Kaibuchi, S. Kuroda, and M. Amano. 1999. Regulation of the Cytoskeleton and Cell Adhesion by the Rho Family GTPases in Mammalian Cells. Annual review of biochemistry. 68(1):459–486, doi: 10.1146/annurev.biochem.68.1.459, https://www.annualreviews.org/doi/abs/10.1146/annurev.biochem.68.1.459.

3. Raftopoulou, M., and A. Hall. 2004. Cell migration: Rho GTPases lead the way. Dev Biol. 265(1):23–32, doi: 10.1016/j.ydbio.2003.06.003, https://www.ncbi.nlm.nih.gov/pubmed/14697350.

4. Lu, S., H. Jang, S. Muratcioglu, A. Gursoy, O. Keskin, R. Nussinov, and J. Zhang. 2016. Ras Conformational Ensembles, Allostery, and Signaling. Chemical Reviews. 116(11):6607–6665, doi: 10.1021/acs.chemrev.5b00542, https://doi.org/10.1021/acs.chemrev.5b00542.

5. Wennerberg, K., K. L. Rossman, and C. J. Der. 2005. The Ras superfamily at a glance. Journal of Cell Science. 118(5):843–846, doi: 10.1242/jcs.01660, https://doi.org/10.1242/jcs.01660.

6. Nethe, M., and P. L. Hordijk. 2010. The role of ubiquitylation and degradation in RhoGTPase signalling. J Cell Sci. 123(Pt 23):4011–4018, doi: 10.1242/jcs.078360, https://www.ncbi.nlm.nih.gov/pubmed/21084561 (https://doi.org/10.1242/jcs.078360).

7. Kumawat, A., S. Chakrabarty, and K. Kulkarni. 2017. Nucleotide Dependent Switching in Rho GTPase: Conformational Heterogeneity and Competing Molecular Interactions. Sci Rep. 7(1):45829, doi: 10.1038/srep45829, https://www.ncbi.nlm.nih.gov/pubmed/28374773.

8. Gorfe, A. A., B. J. Grant, and J. A. McCammon. 2008. Mapping the Nucleotide and Isoform-Dependent Structural and Dynamical Features of Ras Proteins. Structure. 16(6):885–896, doi: https://doi.org/10.1016/j.str.2008.03.009, https://www.sciencedirect.com/science/article/pii/S0969212608001494.

9. Vetter, I. R., and A. Wittinghofer. 2001. The Guanine Nucleotide-Binding Switch in Three Dimensions. Science. 294(5545):1299–1304, doi: doi:10.1126/science.1062023, https://www.science.org/doi/abs/10.1126/science.1062023.

10. Cherfils, J., and M. Zeghouf. 2013. Regulation of small GTPases by GEFs, GAPs, and GDIs. Physiol Rev. 93(1):269–309, doi: 10.1152/physrev.00003.2012, https://www.ncbi.nlm.nih.gov/pubmed/23303910 (https://doi.org/10.1152/physrev.00003.2012).

11. Lowy, D. R., and B. M. Willumsen. 1993. Function and regulation of ras. Annual review of biochemistry. 62(1):851–891, doi: 10.1146/annurev.bi.62.070193.004223, https://www.ncbi.nlm.nih.gov/pubmed/8352603.

12. Hall, A. 2012. Rho family GTPases. Biochem Soc Trans. 40(6):1378–1382, doi: 10.1042/BST20120103, https://www.ncbi.nlm.nih.gov/pubmed/23176484.

13. Etienne-Manneville, S., and A. Hall. 2002. Rho GTPases in cell biology. Nature. 420(6916):629–635, doi: 10.1038/nature01148, https://www.ncbi.nlm.nih.gov/pubmed/12478284 (https://doi.org/10.1038/nature01148).

14. DerMardirossian, C., and G. M. Bokoch. 2005. GDIs: central regulatory molecules in Rho GTPase activation. Trends in cell biology. 15(7):356–363, doi: 10.1016/j.tcb.2005.05.001, https://www.ncbi.nlm.nih.gov/pubmed/15921909 (https://doi.org/10.1016/j.tcb.2005.05.001).

15. Falkenberg, C. V., and L. M. Loew. 2013. Computational analysis of Rho GTPase cycling. PLoS Comput Biol. 9(1):e1002831, doi: 10.1371/journal.pcbi.1002831, https://www.ncbi.nlm.nih.gov/pubmed/23326220 (https://doi.org/10.1371/journal.pcbi.1002831).

16. Cho, H. J., J.-T. Kim, K. E. Baek, B.-Y. Kim, and H. G. Lee. 2019. Regulation of Rho GTPases by RhoGDIs in Human Cancers. Cells. 8(9):1037, https://www.mdpi.com/2073-4409/8/9/1037.

17. Garcia-Mata, R., E. Boulter, and K. Burridge. 2011. The ‘invisible hand’: regulation of RHO GTPases by RHOGDIs. Nature reviews. Molecular cell biology. 12(8):493–504, doi: 10.1038/nrm3153, https://www.ncbi.nlm.nih.gov/pubmed/21779026.

18. Cox, A. D., and C. J. Der. 1992. Protein prenylation: more than just glue? Curr Opin Cell Biol. 4(6):1008–1016, doi: 10.1016/0955-0674(92)90133-w, https://www.ncbi.nlm.nih.gov/pubmed/1485954.

19. Michaelson, D., W. Ali, V. K. Chiu, M. Bergo, J. Silletti, L. Wright, S. G. Young, and M. Philips. 2005. Postprenylation CAAX processing is required for proper localization of Ras but not Rho GTPases. Molecular biology of the cell. 16(4):1606–1616, doi: 10.1091/mbc.e04-11-0960, https://www.ncbi.nlm.nih.gov/pubmed/15659645.

20. Winter-Vann, A. M., and P. J. Casey. 2005. Post-prenylation-processing enzymes as new targets in oncogenesis. Nature reviews. Cancer. 5(5):405–412, doi: 10.1038/nrc1612, https://www.ncbi.nlm.nih.gov/pubmed/15864282.

21. Scheffzek, K., I. Stephan, O. N. Jensen, D. Illenberger, and P. Gierschik. 2000. The Rac-RhoGDI complex and the structural basis for the regulation of Rho proteins by RhoGDI. Nature structural biology. 7(2):122–126, doi: 10.1038/72392, https://www.ncbi.nlm.nih.gov/pubmed/10655614.

22. Hoffman, G. R., N. Nassar, and R. A. Cerione. 2000. Structure of the Rho family GTP-binding protein Cdc42 in complex with the multifunctional regulator RhoGDI. Cell. 100(3):345–356, doi: 10.1016/s0092-8674(00)80670-4, https://www.ncbi.nlm.nih.gov/pubmed/10676816.

23. Grizot, S., J. Faure, F. Fieschi, P. V. Vignais, M. C. Dagher, and E. Pebay-Peyroula. 2001. Crystal structure of the Rac1-RhoGDI complex involved in nadph oxidase activation. Biochemistry. 40(34):10007–10013, doi: 10.1021/bi010288k, https://www.ncbi.nlm.nih.gov/pubmed/11513578.

24. Li, R., and Y. Zheng. 1997. Residues of the Rho family GTPases Rho and Cdc42 that specify sensitivity to Dbl-like guanine nucleotide exchange factors. The Journal of biological chemistry. 272(8):4671–4679, doi: 10.1074/jbc.272.8.4671, https://www.ncbi.nlm.nih.gov/pubmed/9030518.

25. Qiao, J., O. Holian, B. S. Lee, F. Huang, J. Zhang, and H. Lum. 2008. Phosphorylation of GTP dissociation inhibitor by PKA negatively regulates RhoA. Am J Physiol Cell Physiol. 295(5):C1161–1168, doi: 10.1152/ajpcell.00139.2008, https://www.ncbi.nlm.nih.gov/pubmed/18768928.

26. DerMardirossian, C., A. Schnelzer, and G. M. Bokoch. 2004. Phosphorylation of RhoGDI by Pak1 mediates dissociation of Rac GTPase. Molecular cell. 15(1):117–127, doi: 10.1016/j.molcel.2004.05.019, https://www.ncbi.nlm.nih.gov/pubmed/15225553 (https://doi.org/10.1016/j.molcel.2004.05.019).

27. DerMardirossian, C., G. Rocklin, J. Y. Seo, and G. M. Bokoch. 2006. Phosphorylation of RhoGDI by Src regulates Rho GTPase binding and cytosol-membrane cycling. Molecular biology of the cell. 17(11):4760–4768, doi: 10.1091/mbc.e06-06-0533, https://www.ncbi.nlm.nih.gov/pubmed/16943322.

28. Dovas, A., Y. Choi, A. Yoneda, H. A. B. Multhaupt, S.-H. Kwon, D. Kang, E.-S. Oh, and J. R. Couchman. 2010. Serine 34 Phosphorylation of Rho Guanine Dissociation Inhibitor (RhoGDIupalpha) Links Signaling from Conventional Protein Kinase C to RhoGTPase in Cell Adhesion. Journal of Biological Chemistry. 285(30):23296–23308, https://doi.org/10.1074/jbc.m109.098129 (https://doi.org/10.1074/jbc.m109.098129).

29. Knezevic, N., A. Roy, B. Timblin, M. Konstantoulaki, T. Sharma, A. B. Malik, and D. Mehta. 2007. GDI-1 phosphorylation switch at serine 96 induces RhoA activation and increased endothelial permeability. Molecular and cellular biology. 27(18):6323–6333, doi: 10.1128/MCB.00523-07, https://www.ncbi.nlm.nih.gov/pubmed/17636025.

30. Gandhi, P. N., R. M. Gibson, X. Tong, J. Miyoshi, Y. Takai, M. Konieczkowski, J. R. Sedor, and A. L. Wilson-Delfosse. 2004. An activating mutant of Rac1 that fails to interact with Rho GDP-dissociation inhibitor stimulates membrane ruffling in mammalian cells. The Biochemical journal. 378(Pt 2):409–419, doi: 10.1042/BJ20030979, https://www.ncbi.nlm.nih.gov/pubmed/14629200 (https://doi.org/10.1042/bj20030979).

31. Gibson, R. M., P. N. Gandhi, X. Tong, J. Miyoshi, Y. Takai, M. Konieczkowski, J. R. Sedor, and A. L. Wilson-Delfosse. 2004. An activating mutant of Cdc42 that fails to interact with Rho GDP-dissociation inhibitor localizes to the plasma membrane and mediates actin reorganization. Exp Cell Res. 301(2):211–222, doi: 10.1016/j.yexcr.2004.07.033, https://www.ncbi.nlm.nih.gov/pubmed/15530857 (https://doi.org/10.1016/j.yexcr.2004.07.033).

32. Gibson, R. M., and A. L. Wilson-Delfosse. 2001. RhoGDI-binding-defective mutant of Cdc42Hs targets to membranes and activates filopodia formation but does not cycle with the cytosol of mammalian cells. Biochemical Journal. 359(2):285–294, https://doi.org/10.1042/bj3590285 (https://doi.org/10.1042/bj3590285).

33. Golovanov, A. P., T. H. Chuang, C. DerMardirossian, I. Barsukov, D. Hawkins, R. Badii, G. M. Bokoch, L. Y. Lian, and G. C. Roberts. 2001. Structure-activity relationships in flexible protein domains: regulation of rho GTPases by RhoGDI and D4 GDI. J Mol Biol. 305(1):121–135, doi: 10.1006/jmbi.2000.4262, https://www.ncbi.nlm.nih.gov/pubmed/11114252.

34. Lian, L. Y., I. Barsukov, A. P. Golovanov, D. I. Hawkins, R. Badii, K. H. Sze, N. H. Keep, G. M. Bokoch, and G. C. Roberts. 2000. Mapping the binding site for the GTP-binding protein Rac-1 on its inhibitor RhoGDI-1. Structure. 8(1):47–55, doi: 10.1016/s0969-2126(00)00080-0, https://www.ncbi.nlm.nih.gov/pubmed/10673424.

35. Gosser, Y. Q., T. K. Nomanbhoy, B. Aghazadeh, D. Manor, C. Combs, R. A. Cerione, and M. K. Rosen. 1997. C-terminal binding domain of Rho GDP-dissociation inhibitor directs N-terminal inhibitory peptide to GTPases. Nature. 387(6635):814–819, doi: 10.1038/42961, https://www.ncbi.nlm.nih.gov/pubmed/9194563.

36. Ueyama, T., J. Son, T. Kobayashi, T. Hamada, T. Nakamura, H. Sakaguchi, T. Shirafuji, and N. Saito. 2013. Negative Charges in the Flexible N-Terminal Domain of Rho GDP-Dissociation Inhibitors (RhoGDIs) Regulate the Targeting of the RhoGDItextendashRac1 Complex to Membranes. The Journal of Immunology. 191(5):2560–2569, https://doi.org/10.4049/jimmunol.1300209 https://doi.org/10.4049/jimmunol.1300209).

37. Keep, N. H., M. Barnes, I. Barsukov, R. Badii, L. Y. Lian, A. W. Segal, P. C. Moody, and G. C. Roberts. 1997. A modulator of rho family G proteins, rhoGDI, binds these G proteins via an immunoglobulin-like domain and a flexible N-terminal arm. Structure. 5(5):623–633, doi: 10.1016/s0969-2126(97)00218-9, https://www.ncbi.nlm.nih.gov/pubmed/9195882 (https://doi.org/10.1016/s0969-2126(97)00218-9).

38. Golovanov, A. P., D. Hawkins, I. Barsukov, R. Badii, G. M. Bokoch, L. Y. Lian, and G. C. Roberts. 2001. Structural consequences of site-directed mutagenesis in flexible protein domains: NMR characterization of the L(55,56)S mutant of RhoGDI. Eur J Biochem. 268(8):2253–2260, doi: 10.1046/j.1432-1327.2001.02100.x, https://www.ncbi.nlm.nih.gov/pubmed/11298742.

39. Tnimov, Z., Z. Guo, Y. Gambin, U. T. Nguyen, Y. W. Wu, D. Abankwa, A. Stigter, B. M. Collins, H. Waldmann, R. S. Goody, and K. Alexandrov. 2012. Quantitative analysis of prenylated RhoA interaction with its chaperone, RhoGDI. The Journal of biological chemistry. 287(32):26549–26562, doi: 10.1074/jbc.M112.371294, https://www.ncbi.nlm.nih.gov/pubmed/22628549 (https://doi.org/10.1074/jbc.m112.371294).

40. Schrodinger, LLC (2015). The PyMOL Molecular Graphics System, Version 1.8.

41. Eastman, P., J. Swails, J. D. Chodera, R. T. McGibbon, Y. Zhao, K. A. Beauchamp, L. P. Wang, A. C. Simmonett, M. P. Harrigan, C. D. Stern, R. P. Wiewiora, B. R. Brooks, and V. S. Pande. 2017. OpenMM 7: Rapid development of high performance algorithms for molecular dynamics. PLoS Comput Biol. 13(7):e1005659, doi: 10.1371/journal.pcbi.1005659, https://www.ncbi.nlm.nih.gov/pubmed/28746339 (https://doi.org/10.1371/journal.pcbi.1005659).

42. Abraham, M. J., T. Murtola, R. Schulz, S. Páll, J. C. Smith, B. Hess, and E. Lindahl. 2015. GROMACS: High performance molecular simulations through multi-level parallelism from laptops to supercomputers. SoftwareX. 1–2:19-25, https://doi.org/10.1016/j.softx.2015.06.001 (https://doi.org/10.1016/j.softx.2015.06.001).

43. MacKerell, A. D., D. Bashford, M. Bellott, R. L. Dunbrack, J. D. Evanseck, M. J. Field, S. Fischer, J. Gao, H. Guo, S. Ha, D. Joseph-McCarthy, L. Kuchnir, K. Kuczera, F. T. Lau, C. Mattos, S. Michnick, T. Ngo, D. T. Nguyen, B. Prodhom, W. E. Reiher, B. Roux, M. Schlenkrich, J. C. Smith, R. Stote, J. Straub, M. Watanabe, J. Wiorkiewicz-Kuczera, D. Yin, and M. Karplus. 1998. All-atom empirical potential for molecular modeling and dynamics studies of proteins. The journal of physical chemistry. B. 102(18):3586–3616, doi: 10.1021/jp973084f, https://www.ncbi.nlm.nih.gov/pubmed/24889800.

44. Huang, J., and A. D. MacKerell, Jr. 2013. CHARMM36 all-atom additive protein force field: validation based on comparison to NMR data. J Comput Chem. 34(25):2135–2145, doi: 10.1002/jcc.23354, https://www.ncbi.nlm.nih.gov/pubmed/23832629.

45. Bussi, G., D. Donadio, and M. Parrinello. 2007. Canonical sampling through velocity rescaling. J Chem Phys. 126(1):014101, doi: 10.1063/1.2408420, https://www.ncbi.nlm.nih.gov/pubmed/17212484.

46. Parrinello, M., and A. Rahman. 1981. Polymorphic transitions in single crystals: A new molecular dynamics method. J Appl Phys. 52(12):7182–7190, doi: 10.1063/1.328693, <Go to ISI>://WOS:A1981MT07800024.

47. Essmann, U., L. Perera, M. L. Berkowitz, T. Darden, H. Lee, and L. G. Pedersen. 1995. A smooth particle mesh Ewald method. The Journal of Chemical Physics. 103(19):8577–8593, doi: 10.1063/1.470117, <Go to ISI>://WOS:A1995TE36400026.

48. Hess, B., H. Bekker, H. J. C. Berendsen, and J. G. E. M. Fraaije. 1997. LINCS: A linear constraint solver for molecular simulations. Journal of Computational Chemistry. 18(12):1463–1472, doi: 10.1002/(sici)1096-987x(199709)18:12<1463::Aid-jcc4>3.0.Co;2-h, <Go to ISI>://WOS:A1997XT81100004.

49. Kumawat, A., and S. Chakrabarty. 2017. Hidden electrostatic basis of dynamic allostery in a PDZ domain. Proc Natl Acad Sci U S A. 114(29):E5825–E5834, doi: 10.1073/pnas.1705311114, https://www.ncbi.nlm.nih.gov/pubmed/28634294.

50. Liu, J., and R. Nussinov. 2017. Energetic redistribution in allostery to execute protein function. Proc Natl Acad Sci U S A. 114(29):7480–7482, doi: 10.1073/pnas.1709071114, https://www.ncbi.nlm.nih.gov/pubmed/28696318.

51. Kumari, R., R. Kumar, C. Open Source Drug Discovery, and A. Lynn. 2014. g_mmpbsa--a GROMACS tool for high-throughput MM-PBSA calculations. J Chem Inf Model. 54(7):1951–1962, doi: 10.1021/ci500020m, https://www.ncbi.nlm.nih.gov/pubmed/24850022.

52. Tribello, G. A., M. Bonomi, D. Branduardi, C. Camilloni, and G. Bussi. 2014. PLUMED 2: New feathers for an old bird. Computer Physics Communications. 185(2):604–613, doi: 10.1016/j.cpc.2013.09.018, <Go to ISI>://WOS:000329537500020.

53. Kumawat, A., and S. Chakrabarty. 2020. Protonation-Induced Dynamic Allostery in PDZ Domain: Evidence of Perturbation-Independent Universal Response Network. The Journal of Physical Chemistry Letters. 11(21):9026–9031, doi: 10.1021/acs.jpclett.0c02885, https://doi.org/10.1021/acs.jpclett.0c02885.

54. Ellerbroek, S. M., K. Wennerberg, and K. Burridge. 2003. Serine phosphorylation negatively regulates RhoA in vivo. The Journal of biological chemistry. 278(21):19023–19031, doi: 10.1074/jbc.M213066200, https://www.ncbi.nlm.nih.gov/pubmed/12654918.

