## Supporting Information for "Molecular Mechanism of Regulation of RhoA GTPase by Phosphorylation of RhoGDI"

**Table S1:** Phosphorylation code based on available experimental data: Phosphorylation or post-translational modification (PTM) site and their respective effects on specific Rho GTPases

| PTM Site | Kinase | Effect |
|---|---|---|
| Ser34 | PKCα | Promotes dissociation of RhoA |
| Ser96 | PKCα | Promotes dissociation of RhoA |
| Ser101 | PAK1 | Promotes dissociation of Rac1 |
| Ser174 | PAK1 | Promotes dissociation of Rac1 |
| Tyr156 | FER | Promotes dissociation of Rac1 |

**Table S2:** Parameter set used for MM-PBSA calculations

| Parameter | Value |
|---|---|
| APBS solver | Linear |
| Temperature | 300K |
| Boundary conditions (bcfl) | sdh |
| Protein dielectric constant (pdie) | 8 |
| Solvent dielectric constant (sdie) | 80 |
| Grid spacing | 0.5 |
| **coarse factor** | **1.7** |

**Table S3:** $\Delta E_{ij}$ values for all residue pairs involved in allosteric network. All values are reported in kcal/mol.

| Pairs | $E_{ij}^{WT}$ | $E_{ij}^{SP}$ | $\Delta E_{ij}$ $(E_{ij}^{SP} - E_{ij}^{WT})$ |
|---|---|---|---|
| $S34^{RhoGDI}$-$R120^{RhoGDI}$ | 0.9 | -115.8 | -116.7 |
| $E22^{RhoGDI}$-$K127^{RhoGDI}$ | -24.1 | -75.0 | -50.9 |
| $K186^{RhoA}$-$E109^{RhoGDI}$ | -11.8 | -50.9 | -39.1 |
| $E22^{RhoGDI}$-$K186^{RhoGDI}$ | -12.0 | -41.1 | -29.1 |
| $R68^{RhoA}$-$R185^{RhoGDI}$ | -70.2 | -97.5 | -27.3 |
| $H105^{RhoA}$-$E22^{RhoGDI}$ | -18.3 | -44.6 | -26.3 |
| $E17^{RhoGDI}$-$K141^{RhoGDI}$ | -46.3 | -62.6 | -16.3 |
| $E17^{RhoGDI}$-$K138^{RhoGDI}$ | -21.9 | -36.5 | -14.6 |
| $S188^{RhoA}$-$E164^{RhoGDI}$ | 1.2 | -3.7 | -4.9 |
| $E19^{RhoGDI}$-$R180^{RhoGDI}$ | -11.4 | -15.9 | -4.5 |
| $D65^{RhoA}$-$K33^{RhoGDI}$ | -63.9 | -67.5 | -3.6 |
| $R68^{RhoA}$-$E337^{RhoGDI}$ | -26.4 | -22.5 | 3.9 |
| $V64^{RhoGDI}$-$R120^{RhoGDI}$ | -30.1 | -23.3 | 6.8 |
| $R68^{RhoA}$-$K33^{RhoGDI}$ | 18.2 | 25.0 | 6.8 |

| | | | |
|---|---|---|---|
| K186$^{RhoA}$-E163$^{RhoGDI}$ | -38.0 | -31.0 | 7.0 |
| CYSG190$^{RhoA}$-K167$^{RhoGDI}$ | -42.0 | -34.1 | 7.9 |
| R182$^{RhoA}$-D21$^{RhoGDI}$ | -16.4 | -7.4 | 9.0 |
| D65$^{RhoA}$-E37$^{RhoGDI}$ | 28.7 | 38.1 | 9.4 |
| K187$^{RhoA}$-E17$^{RhoGDI}$ | -36.6 | -24.9 | 11.7 |
| K186$^{RhoA}$-E19$^{RhoGDI}$ | -24.6 | -8.0 | 16.6 |
| R185$^{RhoA}$-E22$^{RhoGDI}$ | -36.6 | -17.5 | 19.1 |
| R185$^{RhoA}$-D21$^{RhoGDI}$ | -40.1 | -12.9 | 27.2 |
| K104$^{RhoA}$-E22$^{RhoGDI}$ | -46.4 | -17.9 | 28.5 |
| K186$^{RhoA}$-E20$^{RhoGDI}$ | -55.3 | -5.5 | 49.8 |

**Table S4:** Residue pairs with largest hydrogen bond occupancy differences $\Delta Hb_{ij}$

| Pairs | $Hb_{ij}^{WT}$(%) | $Hb_{ij}^{SP}$(%) | $\Delta Hb_{ij}$ $(Hb_{ij}^{SP} - Hb_{ij}^{WT})$(%) |
|---|---|---|---|
| S34$^{RhoGDI}$-R120$^{RhoGDI}$ | 0.0 | 62.0 | 62.0 |
| V67$^{RhoGDI}$-R120$^{RhoGDI}$ | 48.0 | 20.0 | -28.0 |
| R68$^{RhoA}$-D185$^{RhoGDI}$ | 50.0 | 96.0 | 46.0 |
| K104$^{RhoA}$-E22$^{RhoGDI}$ | 27.0 | 1.0 | -26.0 |
| E22$^{RhoGDI}$-K185$^{RhoA}$ | 21.0 | 0.0 | -21.0 |
| E22$^{RhoGDI}$-K127$^{RhoGDI}$ | 2.0 | 48.0 | 46.0 |
| H23$^{RhoGDI}$-D184$^{RhoGDI}$ | 0.0 | 44.0 | 44.0 |
| D21$^{RhoGDI}$-K185$^{RhoA}$ | 20.0 | 0.0 | -20.0 |
| E20$^{RhoGDI}$-K186$^{RhoA}$ | 36.0 | 0.0 | -36.0 |
| E17$^{RhoGDI}$-K186$^{RhoA}$ | 30.0 | 0.0 | -30.0 |
| E17$^{RhoGDI}$-K141$^{RhoGDI}$ | 21.0 | 42.0 | 21.0 |
| E109$^{RhoGDI}$-K186$^{RhoA}$ | 0.0 | 20.0 | 20.0 |
| E163$^{RhoGDI}$-K187$^{RhoA}$ | 7.0 | 70.0 | 63.0 |
| E164$^{RhoGDI}$-S188$^{RhoA}$ | 9.0 | 37.0 | 28.0 |
| K167$^{RhoGDI}$-CYSG$^{RhoA}$ | 61.0 | 15.0 | -46.0 |
| D140$^{RhoGDI}$-K186$^{RhoA}$ | 74.0 | 13.0 | -61 |

| | | | |
|---|---|---|---|
| A8$^{RhoGDI}$-A12$^{RhoGDI}$ | 70.0 | 11.0 | -59.0 |
| N10$^{RhoGDI}$-I14$^{RhoGDI}$ | 65.0 | 8.0 | -57.0 |
| P30$^{RhoGDI}$-R68$^{RhoA}$ | 17.0 | 73.0 | 56.0 |
| M162$^{RhoGDI-}$K185$^{RhoA}$ | 0.0 | 30.0 | 30.0 |
